# Short distance non-autonomy and intercellular transfer of chitin synthase in Drosophila

**DOI:** 10.1101/2020.05.24.113803

**Authors:** Paul N. Adler

## Abstract

The complex structure of insect exoskeleton has inspired material scientists and engineers. Chitin is a major component of the cuticle and it is synthesized by the enzyme chitin synthase. There is a single chitin synthase gene (*kkv*) in Drosophila facilitating research on the function of chitin. Previous editing of *kkv* lead to the recovery of a viable hypomorphic allele. Experiments described in this paper argue that a reduction in chitin synthase leads to the shafts of sensory bristles becoming fragile and frequently breaking off as the animals age. This is likely due to reduced chitin levels and further suggests that chitin plays a role in resilience of insect cuticle. The different layers in cuticle are continuous across the many epidermal cells that secrete the cuticle that covers the body. Little is known about the mechanisms responsible for this continuity. Using genetic mosaics and scanning electron microscopy this paper establishes that *kkv* shows short range cell non-autonomy. It also provides evidence for 2 possible mechanisms. One is the intercellular transfer of Kkv protein from one cell to its neighbors and the second is the deposition of cuticular material across the boundaries of neighboring cells.

## Introduction

### Review of complexity and flexible properties of cuticle

The cuticular exoskeleton of insects and other arthropods play a central role in the life style of these extremely successful groups of animals and it has served as a model for material scientists designing composite materials (Fernandez and Ingber, 2012; Grunenfelder et al., 2014; Rajabi et al., 2015; Vincent and Wegst, 2004). Cuticle is composed of several layers that are qualitatively different from one another (Fig 1) (Moussian et al., 2007; Payre, 2004; Sobala and Adler, 2016). Several different nomenclatures have been used to identify these, and in part this reflects the structural variation in insect cuticles. I will use the term envelope to describe the outermost layer that is lipid rich and serves as a barrier to water loss and aids in hydrophobicity. I use the term epicuticle for the next layer. Both of these are rather thin in the Drosophila cuticles studied in this paper. The innermost layer is the procuticle, which is composed of multiple sub-layers of chitin and protein. The chitin fibrils are arranged as parallel arrays and each layer is rotated with respect to its neighboring layers (Bouligand, 1972; Moussian, 2013; Moussian et al., 2006). The array rotation gives rise to the layered appearance of the procuticle. The procuticle is typically the thickest layer and the arrays of chitin are thought to provide strength and perhaps resilience to failure. In addition to these 3 layers various authors have proposed an additional layer that is juxtaposed between the apical surface of the epidermal cells and the basal region of the procuticle. This fourth layer was given the names such as the assembly zone or adhesion layer (Fristrom et al., 1993; Locke, 1961; Moussian et al., 2005; Schmidt, 1956; Sobala and Adler, 2016). The suggested function can be gleamed from their names. My observations suggest that both of these two types of layers can be found in at least some developing cuticles and that they are different from one another (Sobala and Adler, 2016). The insect cuticle is extremely variable in terms of its physical properties. For example, the Young’s modulus of insect cuticle varies over more than 6 orders of magnitude (Vincent and Wegst, 2004). Qualitative and quantitative differences in the molecular components, their degree of cross linking and the degree of hydration are likely responsible for this variation. To a first approximation, it is generally thought that the layers are synthesized in the order envelope, exocuticle and procuticle but there is likely at least some “maturation” that takes place out of order (Moussian et al., 2006; Sobala and Adler, 2016).

**Figure 1.**
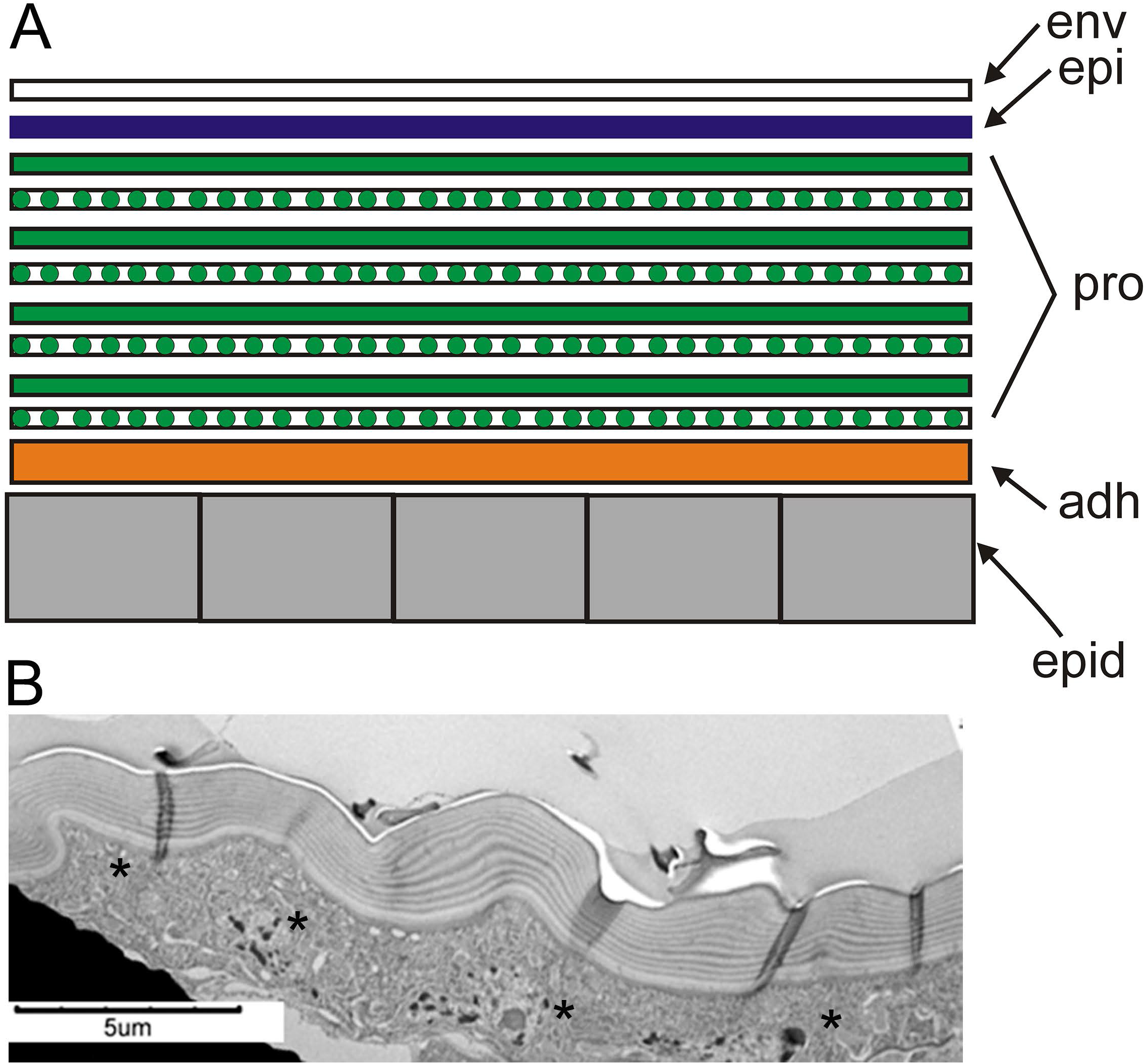
Layered structure of cuticle. A. The cartoon shows the location of the various different layers in cuticle. B. A transmission electron microscope image of abdominal cuticle. Note the very prominent layering in the procuticle and that individual bands are continuous across the image. The asterisks indicate the location of junctional complexes. These are difficult to see at this magnification but the relative position of the junctional complexes could be identified at a higher magnification. The micrograph extends over 3 complete cells and 2 partial cells.

Insect cuticle is a complicated material containing a large number of different components. For example, more than 100 cuticle protein genes are found in the Drosophila genome (Ioannidou et al., 2014; Willis, 2010) and it has been suggested that this may be an underestimate (Sobala and Adler, 2016). During the synthesis of the Drosophila wing cuticle 83 annotated cuticle protein genes are expressed (Sobala and Adler, 2016). In contrast, there is a single gene (*kkv*) that encodes the enzyme (chitin synthase) that synthesizes chitin. That there is a single chitin synthase makes it an attractive target for studying details of cuticle deposition. I recently found that an edited *kkv* which fused an smFP tag to the c terminal end of the protein resulted in a viable hypomorphic allele (Adler, 2020). This was identified on the basis of a phenotype in the hairs that cover the wing. With further studies we have found additional phenotypes associated with this mutation and one of these suggests the cuticle synthesized in mutants is less robust to the wear and tear of life.

### The importance of continuity and lack of discontinuities for robustness

Insect cuticle is not only layered but the layers are continuous across the many epidermal cells in a body region that secrete it (Fig 1B). This continuity is likely important for the robustness of the cuticle as discontinuities could lead to stress concentration and cracks whose propagation could lead to failure (Iremonger and Wood, 1969) (https://en.wikipedia.org/wiki/Stress_concentration). Particularly for the deposition of the procuticle, synchrony seems likely to be crucial as even slight differences in the timing of chitin array deposition between neighboring cells might lead to discontinuities in the chitin arrays.

### How is the continuity achieved?

The synthesis of cuticle is under at least indirect control of ecdysone (Ali et al., 2013; Cui et al., 2009; Gangishetti et al., 2012; Kozlova et al., 2009; Wang et al., 2010). It is not clear however, that a burst of hormone is adequate for ensuring cuticle layer continuity. Neighboring cells are likely to respond to a pulse of ecdysone at a very similar time but the synthesis of cuticle does not start for several days after the formation of white prepupae and it seems possible that if ecdysone pulses were the sole regulatory mechanism that individual cells timing might become unsynchronized enough to result in discontinuities. Several observations establish that the development of cells in a tissue is not always temporally uniform. For example, it has been known for decades that the morphogenesis of hairs (trichomes) on the wing blade is not synchronous across the wing. Hair morphogenesis begins first near the distal tip of the wing and it progresses in a patchy fashion proximally (Wong and Adler, 1993). However, as I report here this temporal patchiness does not appear to be true for the later wing cuticle deposition implying something synchronizes it later. The deposition of cuticle is also not uniformly synchronous across the apical surface of individual Drosophila epidermal cells. For the wing, patches of envelop first appear over developing wing hairs and then later over the wing blade (Sobala and Adler, 2016; Wong and Adler, 1993). Except for the hair/wing blade variation, the patches appear to be distributed randomly on individual cells. Over time, the patches expand to cover the entire apical surface of the epidermal cells. This is accomplished earlier over the hairs than the wing blade region of these cells.

### Timing models

The simplest mechanism to regulate the timing of exoskeleton development would be to have a program initiate due to a pulse of ecdysone and to free run from that point. Such a simple model does not explain the different timing patterns noted above and it is not clear how precise such a mechanism would be. To insure a tighter temporal control for cuticle deposition a signaling/signal transduction system could synchronize cells in different tissues at various stages. Such a system could work by activating one or more transcription factors that resulted in the expression of genes required for cuticle deposition. RNA seq has identified many genes whose expression level changes dramatically during the process of cuticle deposition consistent with such a model (Sobala and Adler, 2016).

It is also possible that a pathway that functioned at the morphogenesis level helps ensure the deposition of cuticle is synchronized between neighboring cells. For example, the movement in the extracellular space or by intercellular transport of proteins that play an important role in cuticle deposition could do that. If this happened it could be detected as cell non-autonomy. Previous experiments that only examined the hairs indicated that *kkv* functioned cell autonomously (Ren et al., 2005). As is described in the results I obtained evidence for both the intercellular movement of Kkv over short distances and evidence for short-range cell non-autonomy.

## Results

### Cuticle deposition is uniform across a tissue unlike wing hair morphogenesis

The morphogenesis of wing hairs begins in the distal part of the wing and moves proximally; however, this is not accomplished by the equivalent of the morphogenetic furrow in the eye disc. Rather the timing is patchy with some neighboring cells being at different stages in hair morphogenesis (Fig 2A) (Lu et al., 2015; Wong and Adler, 1993). As morphogenesis continues the delayed hairs catch up to the earlier forming ones. To determine if the deposition of cuticle followed a similar time course I used a chitin reporter and examined wing cuticle deposition by in vivo imaging (Sobala et al., 2015). At all times examined (42, 44 and 46 hr awp (after white prepupae)) neighboring cells showed similar levels of the reporter (Fig 2C). I carried out similar experiments where we localized Kkv::NG and similarly in all cases neighboring cells always showed similar levels and localization of Kkv (Fig 2B). We also examined pigmentation in developing pupae and once again, neighboring cells showed a similar level of pigmentation (Fig S1). Hence the mechanism used to initiate hair outgrowth appears to be different from the mechanism used to initiate chitin deposition and Kkv synthesis.

**Figure 2.**
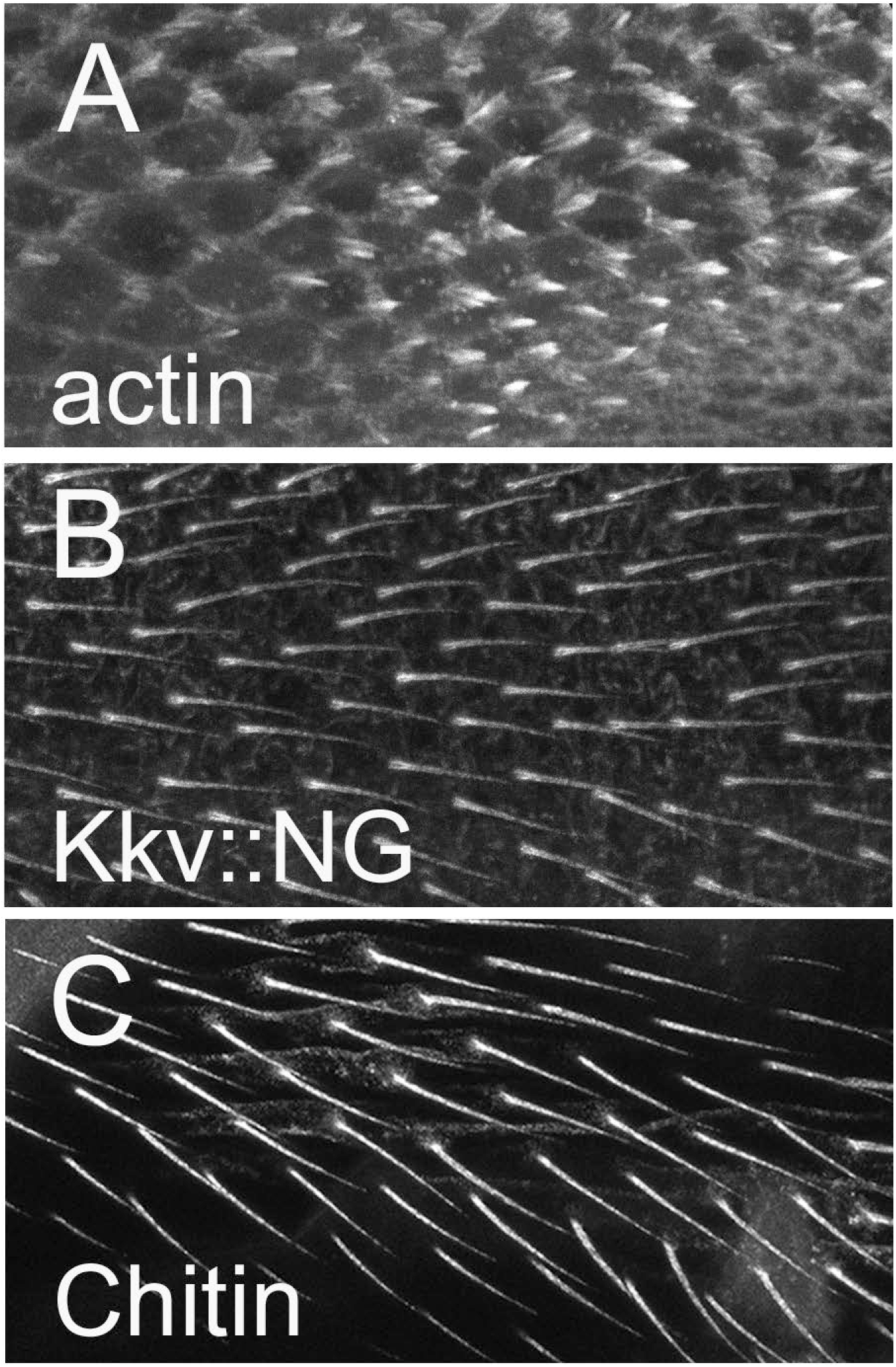
Neighboring cells are synchronous in chitin deposition and Kkv accumulation. A. An in vivo micrograph of actin in a 32 hr wing. Note how the process of hair out growth is not synchronous. Instead, it is quite patchy. B. An in vivo micrograph of Kkv::NG in a 42 Hr pupal wing. Note all of the hairs show a similar level of Kkv::NG accumulation. C. An in vivo micrograph showing the accumulation of chitin (by the use of the ChtVis chitin reporter (Sobala et al., 2015) in a 46 hr pupal wing.

### The cuticle of a hypomorphic allele of *kkv* is less robust than wild type cuticle

As described previously I used Crispr/Cas9 to tag the endogenous *kkv* gene with either Neon Green or smFP (Adler, 2020). Flies homozygous for each of the edits were viable and showed no dramatic phenotype when examined in a stereomicroscope. However, when we mounted wings for examination in the compound microscope we observed thin, bent and generally deformed wing hairs (trichomes) in the smFP homozygotes but not the NG homozygotes (Adler, 2020). We further established that the Kkv::smFP protein accumulated to a much lower level than Kkv::NG. When we examined *kkv∷smFP/Df* flies the hair phenotype appeared slightly stronger. These two observations let me to conclude *kkv∷smFP* was a hypomorph due to the protein not accumulating efficiently (Adler, 2020). I subsequently noticed that a minority of the *kkv∷smFP/Df* flies had deformed wings soon after or at eclosion (Fig 3). This suggested a problem in expansion of the wing or eclosion from the pupal case. We found the frequency of flies showing the phenotype was significantly higher for *kkv∷smFP/Df, kkv*::smFP/*kkv*^*1*^ (*kkv*^*1*^ is listed as an amorphic allele on FyBase (Dos Santos et al., 2015)) and *kkv::smFP* flies compared to Ore-R or *kkv∷NG* flies (Fig 3). We also noticed that some of the *kkv*∷*smFP* containing flies showed a slight downward curve to the wing. While the *kkv* hypomorphic individuals are able to fly, by casual observation, they appeared less active than normal flies.

**Figure 3.**
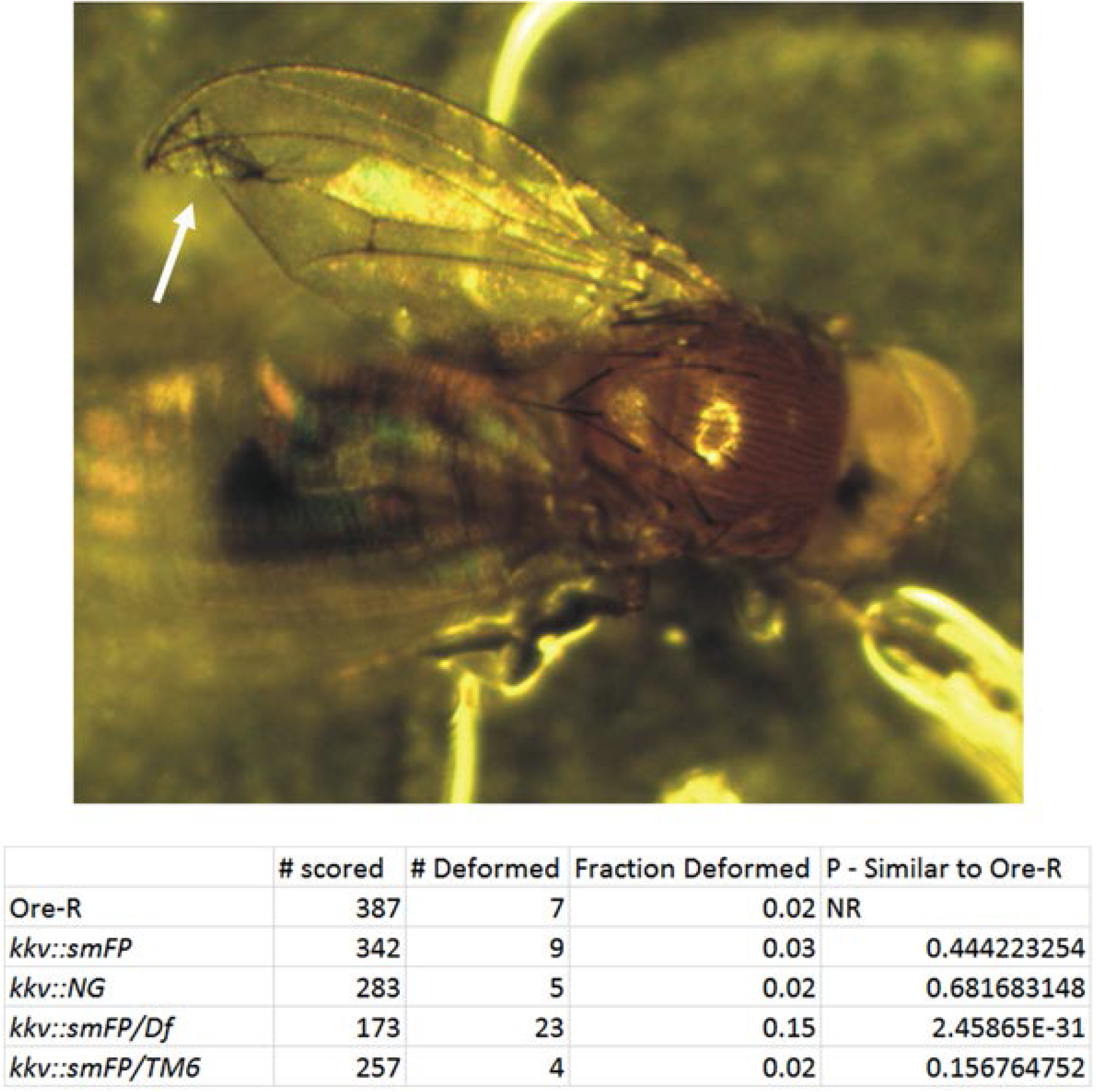
Deformed wings in newly eclosed flies that carry the *kkv∷smFP* allele. The upper panel is a *kkv∷smFP/Df* fly with a folded wing (arrow). The lower panel shows the frequency of this phenotype for several different genotypes and the p values are from a chi-square test of a comparison of the observed frequency to that predicted if the frequency in question was no different from *Ore-R.*

To test the hypothesis that the cuticle synthesized in a hypomorphic mutant was less robust to the wear and tear of life we followed adult flies that were either wt or *kkv* hypomorphs over time to see if defects arose more rapidly in the hypomorphs. We scored 3 phenotypes: life span, the loss of thoracic macrocheatae or wing defects. All of the various mutants were compared to wild type Ore-R flies. To reduce the length of time that these experiments took we kept the adult flies at 29.5 °C or 29.3°C, which shortens lifespan. We carried out 2 separate experiments using slightly different conditions (see methods) and in both cases the Ore-R flies lived longer than the *kkv* edited flies (Fig S2). The data suggested that this was not simply due to decreased Kkv levels as *kkv∷smFP/Df* flies lived slightly longer than the *kkv∷smFP* flies. The basis for the shorter life span is unclear.

A significant difference was seen for the loss of thoracic macrocheatae with aging that did appear to be due to *kkv* levels as the strongest phenotypes were seen for *kkv∷smFP/Df* and *kkv∷smFP/kkv*^*1*^ flies (Fig 4 H-K). Most of those flies that lived for 20 days or longer showed the loss of at least one of those bristles and in most cases multiple bristle were lost. I also observed the loss of microcheatae (Fig 4) but did not quantify this. In all, or at least most cases, the bristle shaft was lost but the socket cell remained (Fig 4G). In addition to the two experiments where 10 females and 5 males were cultured together, we also carried out a small scale experiment where we followed individual female flies for 20 days. This experiment established that the loss was progressive. That is, we usually observed the loss of one or two bristles followed by the subsequent loss of additional bristles (Fig 4DEF). Bristle loss was less frequent but still common in *kkv∷smFP* homozygoes. In contrast, the loss of bristles associated with aging was rarely seen for Ore-r (Fig 4 ABC, H-K).

**Figure 4.**
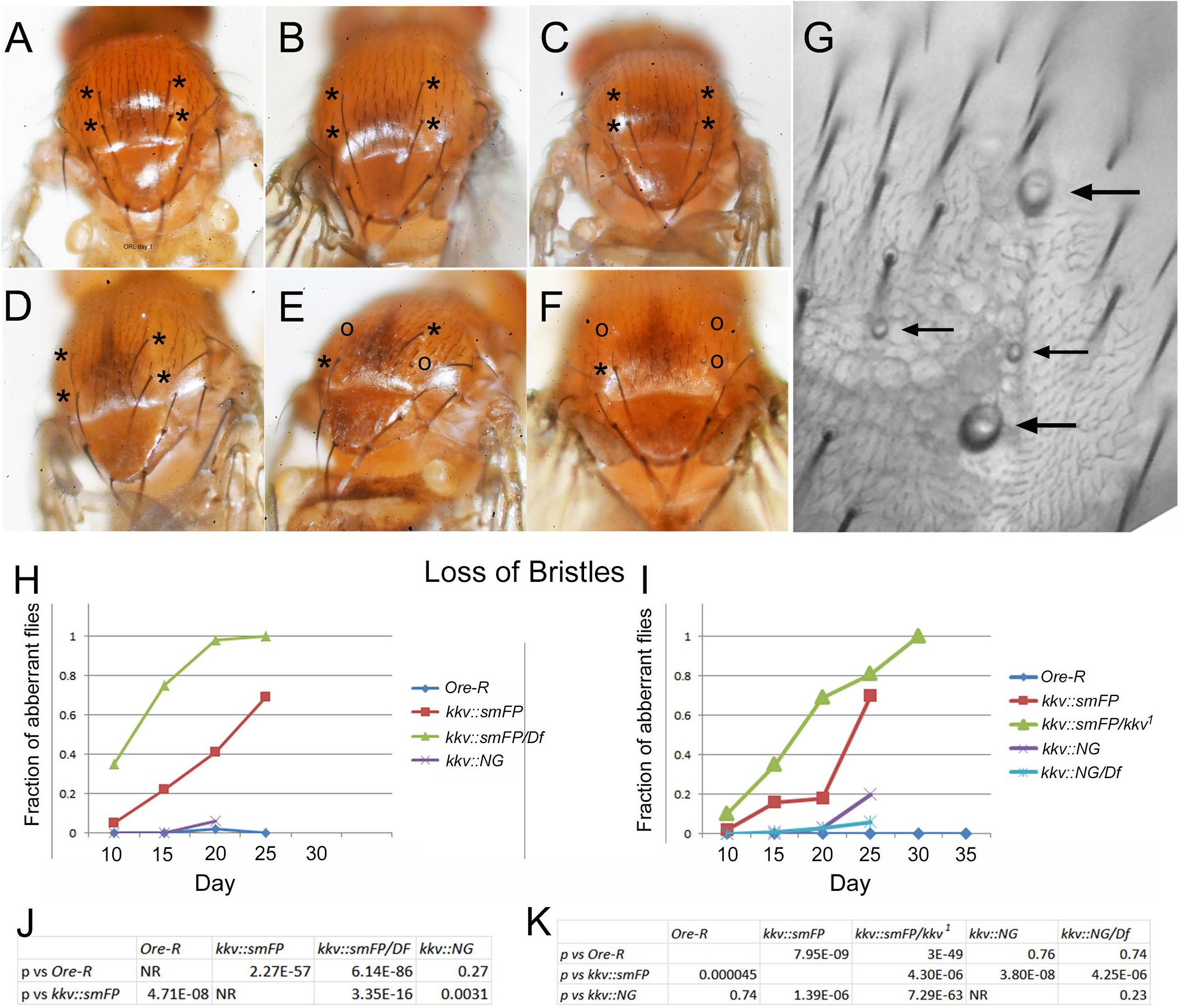
Bristle shaft loss associated with aging of *kkv∷smFP* flies. A. A 6 hr old Ore-R fly. B. The same fly from panel A at 15 days old Ore-R. C. The same fly from panel A at 20 days old. D. A 1 day old *kkv∷smFP/Df* fly. E. The same fly from panel D at 15 days old. F. The same fly from panel D at 20 days old. Asterisks mark the 4 dorsocentral bristles. These are the bristles most commonly lost over time in *kkv∷smFP/Df* flies. O marks dorsocentral bristles shafts that have been lost. In many cases the socket is still visible. G. A *kkv∷snFP/Df* fly mounted and imaged by bright field microscopy. The large arrows point to the remaining socket cell after the shaft cell has been lost. The small arrows point to the socket cells remaining after the shaft of microcheatae were lost. Note the progressive loss of bristles in panels D-F and the lack of bristle shafts in wild type flies (A-C).

We also scored the flies in these experiments for loss of sections of the wing. The losses routinely began at the wing margin. Surprisingly, this turned out to be substantially more common in Ore-R flies than in the flies with edited *kkv* genes (Fig S3). Possible reasons for this are explored in the discussion.

### Evidence for non-autonomy

Published experiments established that in genetic mosaics *kkv* (chitin synthase) acted cell autonomously with regard to wing hairs (Ren et al., 2005). A limitation of those experiments is that during cuticle formation the hairs are located on pedestals in the center of the cell (Mitchell et al., 1990; Sobala and Adler, 2016). Thus, we could have missed short range (i.e. part of a cell) cell non-autonomy. In similar experiments, we observed clones in the abdomen and notum due to their lack of pigmentation and when examined using a dissecting microscope the pigmentation appeared to have sharp borders consistent with cell autonomous function. In the wing pigment appears to accumulate in the apical part of the procuticle (Riedel et al., 2011), so it is not surprising that a loss of chitin arrays in a *kkv* mutant clone results in a lack of pigmentation.

I reinvestigated the cell autonomy of pigmentation in *kkv* mutant clones located in abdomen using mounted cuticle and a compound microscope. I generated flies that carried flip out clones that knocked down the expression of *kkv* (*w hs-flp*; *Ay-Gal4; UAS-kkv-rnai*). When I examined the resulting clones using a high NA (1.3) oil immersion 40X objective it was obvious that the clone boundaries were not sharp. We examined 61 Clones and none showed a sharp pigmentation boundary (Fig S4C-J) It is worth noting that naturally occurring pigmentation boundaries in the abdomen are relatively sharp so in structurally normal cuticle there is nothing that prevents a sharp boundary (Fig S4AB). In similar clones (i.e. *w hs-flp*; *Ay-Gal4; UAS-kkv-rnai*) the phenotype in wing hairs suggested this genotype results in a range of phenotypes that vary from hypomorphic to strong, close to null *kkv* phenotypes.

### Sem of wing clones

In an attempt to assay more directly for non-autonomy of *kkv* mutant clones I needed an assay that allowed us to identify *kkv* clones and examine them at a higher resolution than is possible in the light microscope. I found that I could fracture adult cuticle and by scanning electron microscopy detect a layered structure. The layers were very distinctive in abdominal cuticle (Fig S5) consistent with the robust layering in abdominal procuticle seen by TEM (Fig 1). I also attached wings to studs in a vertical position, fractured them and then imaged the wings by scanning electron microscopy. In wing cuticle I also detected a layered structure that presumably reflects the banding of chitin in the procuticle (Fig 5BC). The layering was less distinctive (and sometimes hard to detect) than in the abdominal cuticle consistent with TEM observations. We next fractured and observed wings carrying *kkv* loss of function clones. We were able to identify mutant clones by the presence of the *kkv* flaccid hair phenotype (Fig 5DE arrows) (Ren et al., 2005). If there was complete cell autonomy we predicted a sharp change in wing cuticle thickness at wt-mutant clone boundaries (Fig 5A). In contrast, if there was a small degree of cell non-autonomy we predicted that we would see a smooth change in cuticle thickness near the edge of clones (Fig 5A). In all of the clones (n=9) we examined there was a smooth transition in cuticle thickness (Fig 5DE). This transition zone appeared to be restricted to a mutant cell and its direct neighbor.

**Figure 5.**
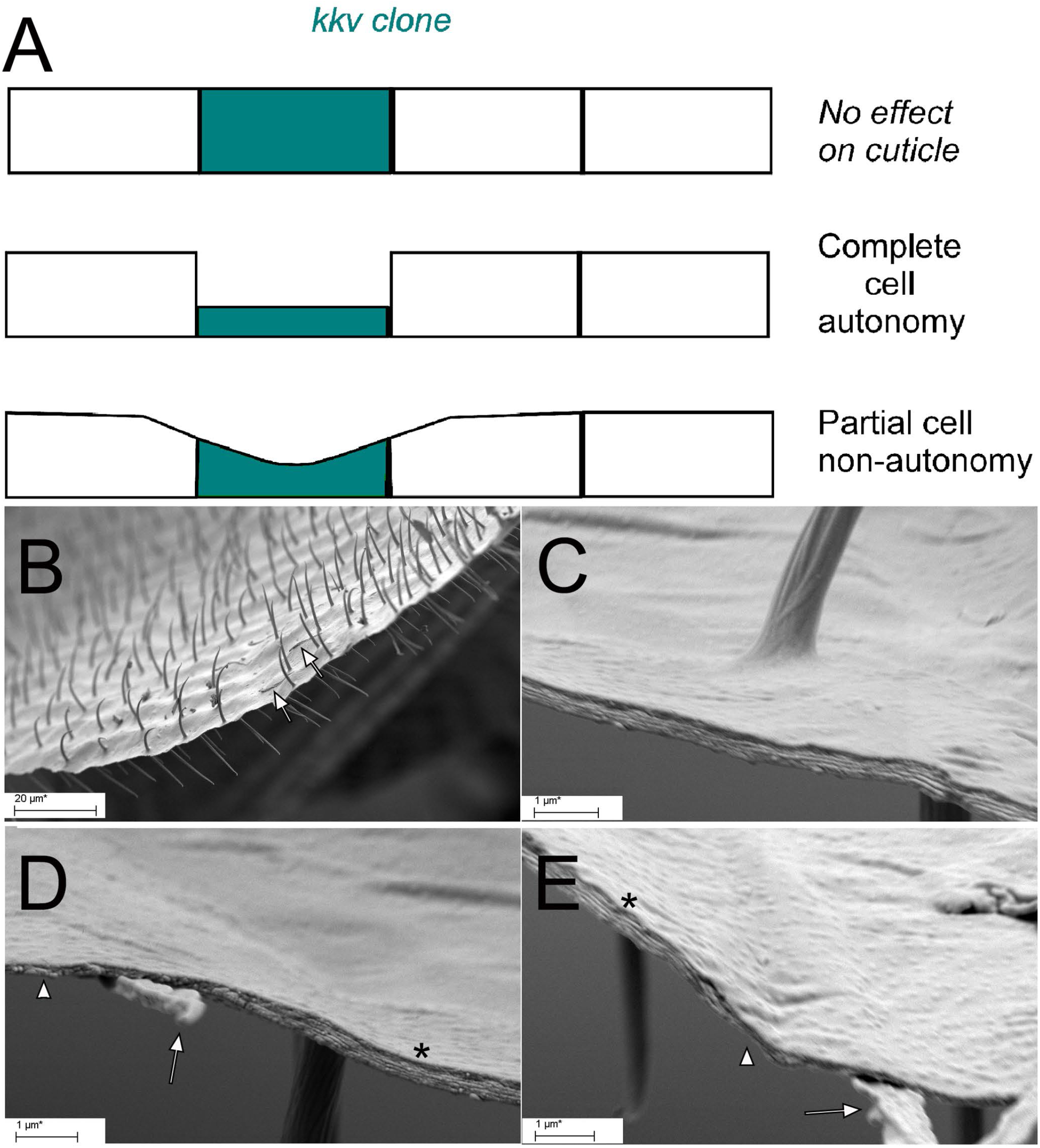
Examination of *kkv* mutant clones by SEM. A. Three possible models for the phenotypic effects of a *kkv* mutant cell (in green) on it and its neighbors. In the uppermost the mutant phenotype is rescued by neighboring wild type cells. In the middle panel the inability to synthesize chitin results in a thinner wing cuticle and this is hypothesized to be entirely cell autonomous. In the lowest most panel there is partial rescue of the ability to synthesize chitin and this is graded from neighboring to mutant cell. B. A relatively low magnification SEM image of a fractured adult wing that contains a clone of *kkv* mutant cells. Arrows point to two of these. C. A region of a wing without any *kkv* clones. Note the relatively consistent cuticle thickness and the parallel layers. D. A region of a wing that contains a mutant cell (mutant hair marked by an arrow). Note how thin the mutant cuticle is (arrowhead) compared to the neighboring wild type cuticle (asterisk). Note also that there is a smooth and graded transition between the mutant and neighboring cell cuticle. E. Another example of a *kkv* mutant clone as in D.

### Possible mechanism for the short-range non-cell autonomy of *kkv*

#### Kkv is shed

In experiments where we imaged Kkv::NG in living pupae we noticed fluorescent puncta in the extracellular space between the pupal cuticle and the epidermal cells that were in the process of synthesizing the adult cuticle (Fig. 6BC). To investigate this in more detail we obtained large Z stacks that extended from the pupal cuticle to below the apical surface of the epidermal cells. Pupal cuticle in 50 hr pupae shows substantial autoflourescence so we examined and compared *Ore-R* and *kkv∷NG* pupae to determine what if any fluorescence was due to the presence of the Kkv-NG protein. The autofluorescence of the thoracic pupal cuticle of Ore-R was spatially relatively even (Fig 6D). In contrast, the fluorescence of pupal cuticle of *kkv-NG* flies was much more uneven with both puncta (arrow) and lines (arrowhead) of bright fluorescence (Fig. 6A). No fluorescence was observed in the region between the pupal cuticle and the apical surface of the epithelial cells in Ore-R pupae (Fig. 6EF). In contrast, in this region many fluorescent puncta were observed in *kkv∷NG* pupae (Fig. 6BC, arrows). A majority of these were located close to the pupal cuticle but some were observed throughout the region. Most of the puncta located close to the pupal cuticle were stable but many of those located lower were mobile (movie S1). Since the fluorescent puncta were only seen when the *kkv∷NG* gene was present we interpret the puncta as evidence of shed Kkv::NG. Since the puncta were located above the impermeable adult cuticle (which is in the process of being synthesized), it seems likely that the Kkv::NG was shed during or after the synthesis of the pupal cuticle and before the synthesis of the adult cuticle began. Consistent with this hypothesis we observed puncta in 28hr pupae, well before the start of adult cuticle deposition (Sobala and Adler, 2016). We also observed puncta in very young pupae (20 hr awp) prior to the detachment of the epithelial cells from the pupal cuticle (Fig. S6B). We observed similar puncta when we examined *ap*>*kkv::NG* pupae (Fig S6C) but not from *ap*>*kkv-R896K*::NG pupae (this mutant protein does not localize correctly to the apical surface (Adler, 2020)) (Fig S6D), suggesting that to be shed Kkv needs to be localized apically.

**Figure 6.**
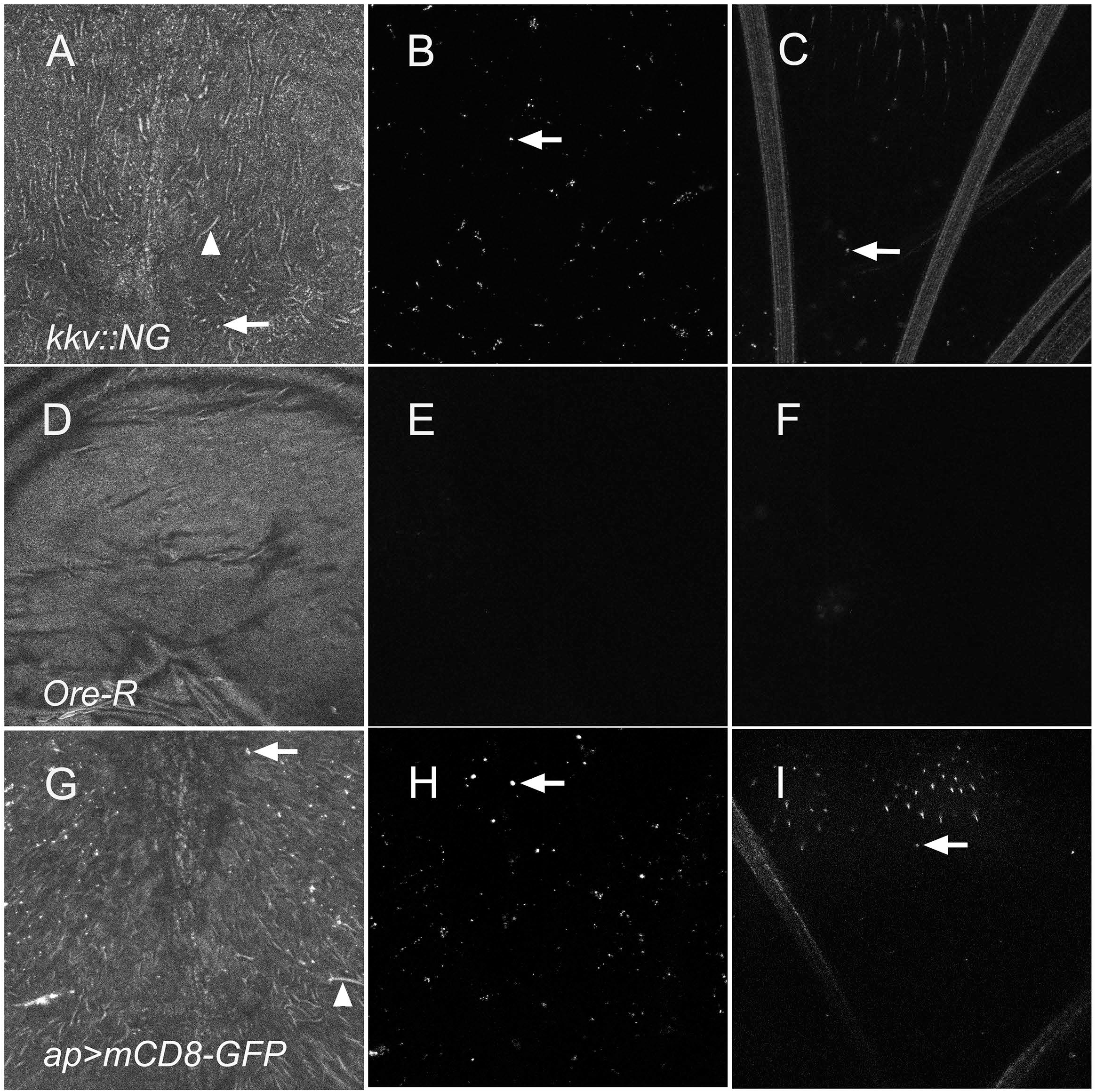
Expression of *kkv∷NG* leads to fluorescent puncta in the space between the pupal cuticle and the apical surface of the notum epithelial cells. A, B, C. *kkv∷NG* pupae. D, E, F. Oregon R pupae. G, H, I. *ap-Ga4; UAS-mCD8*-GFP. A, D and G are maximal projections of the optical stack regions that encompass the notum pupal cuticle. The arrow in A and G points to a likely puncta and the arrowheads to the patterned brightness seen in the pupal cuticle of animals that express a fluorescent membrane protein. B, E and H are maximal projections of the optical stack regions that encompass the region between the notum pupal cuticle and the upper most region of the notum epithelial cells (i.e. the bristles that extend apically). Since there was no fluorescence in E we took the conservative approach of extending this region of the stack by 50%. The arrows in B and H point to one of the many fluorescent puncta. C, F and I are maximal projections of the optical stack regions that encompass the region of cellular material including bristles, hairs and the apical surface of the notum cells. The arrows points to fluorescent puncta.

The highest concentration of puncta were over the dorsal thoracic midline (Fig S6ABC). There were also a large number of puncta over the dorsal abdomen and they were seen at a lower frequency in the wing, legs and head. In the abdomen the puncta tended to align parallel to the segment boundary.

All of the experiments where we detected puncta in living pupae required imaging of the Kkv::NG fusion protein. Experiments described earlier established that NG was a valid reporter for Kkv in bristles so it seemed likely that it was also a valid reporter for Kkv in puncta (Adler, 2020). To test this hypothesis I immunostained pupal cuticle using both anti-NG and anti-Kkv-M antibodies. Among the puncta detected 69.7% stained with both antibodies indicating most puncta contained both NG and Kkv (Fig S7, arrows) supporting the idea that NG is an valid reporter for shed Kkv::NG.

The shedding of Kkv::NG could be specific for Kkv or it could reflect a process that leads to the shedding of many if not all of the proteins located in the apical plasma membrane. To distinguish between these two possibilities we examined live *ap-Gal4*/+; *UAS-mCD8-GFP*/+ pupae. These animals showed a large number of fluorescent puncta present in the space between the pupal cuticle and the apical surface of the epithelial cells (Fig 6GHI). As was the case for the puncta in *kkv∷NG* pupae many of the puncta were mobile. I concluded that the shedding of membrane proteins is not specific for Kkv or proteins involved in cuticle deposition.

The pupal cuticle of Drosophila appears relatively transparent and uniform in bright field optics. However, in carrying out these in vivo imaging experiments we observed that the autofluorescence of the thoracic and abdominal pupal cuticles was quite distinct (Fig S8). The autofluorescence of the thoracic pupal cuticle was splotchy but without distinctive morphology. In contrast, the autofluorescence of the abdominal pupal cuticle showed a pattern of bright elongated lines. Our first thought when observing this was that the bright lines represented cell boundaries however attempts to establish this were unsuccessful.

### Flip out clones - Transfer of Kkv::NG puncta

The observation that Kkv::NG was shed raised the possibility that this could be a mechanism to provide for cell non-autonomy of *kkv.* One possible mechanism is the apical secretion of Kkv and its subsequent lateral movement prior to it synthesizing chitin. An alternative possibility is the intercellular movement of Kkv to neighboring cells followed by secretion or apical localization and chitin synthesis. In an attempt to get evidence for either of these mechanisms we generated flip out clones comprised largely of single cells that expressed Kkv::NG and then looked by in vivo confocal microscopy for lateral movement of Kkv::NG beyond the clone cell. We observed 53 such clones during the deposition of the procuticle and for 41 of these we observed Kkv::NG puncta beyond the lateral edge of the clone cell (Fig 7AB). These puncta could be localized apically over the neighboring cells or in the neighboring cell. From Z section series, I was able to determine whether the puncta were at or above the apical surface of neighboring cells, not at or above the apical surface but in the apical most half of the neighboring cell or in the basal most half. I scored 154 Kkv::NG puncta and found 26 that mapped apical to the neighboring cell, 74 that mapped to the apical half of the neighboring cell and 54 that mapped in the basal half of the neighboring cell. Thus, most of the puncta were not located apical to the clone cell, which is where we observed shed Kkv::NG in earlier experiments on animals after the completion of pupal cuticle synthesis suggesting a different process. Rather, most by their location appeared to be inside of neighboring cells. One possible explanation for these observations is the intercellular transfer of Kkv::NG (or its mRNA) to neighboring cells with some of that protein subsequently being secreted apically. A second is the apical secretion of the Kkv::NG puncta by the clone cell with the endocytosis of the puncta by neighboring cells. In either case Kkv::NG appears to spread to neighboring cells. I did not observe obvious cases where the puncta moved multiple cells but since the neighboring cells were not labelled I could not rule out a small number of puncta moving more than one cell laterally.

**Figure 7.**
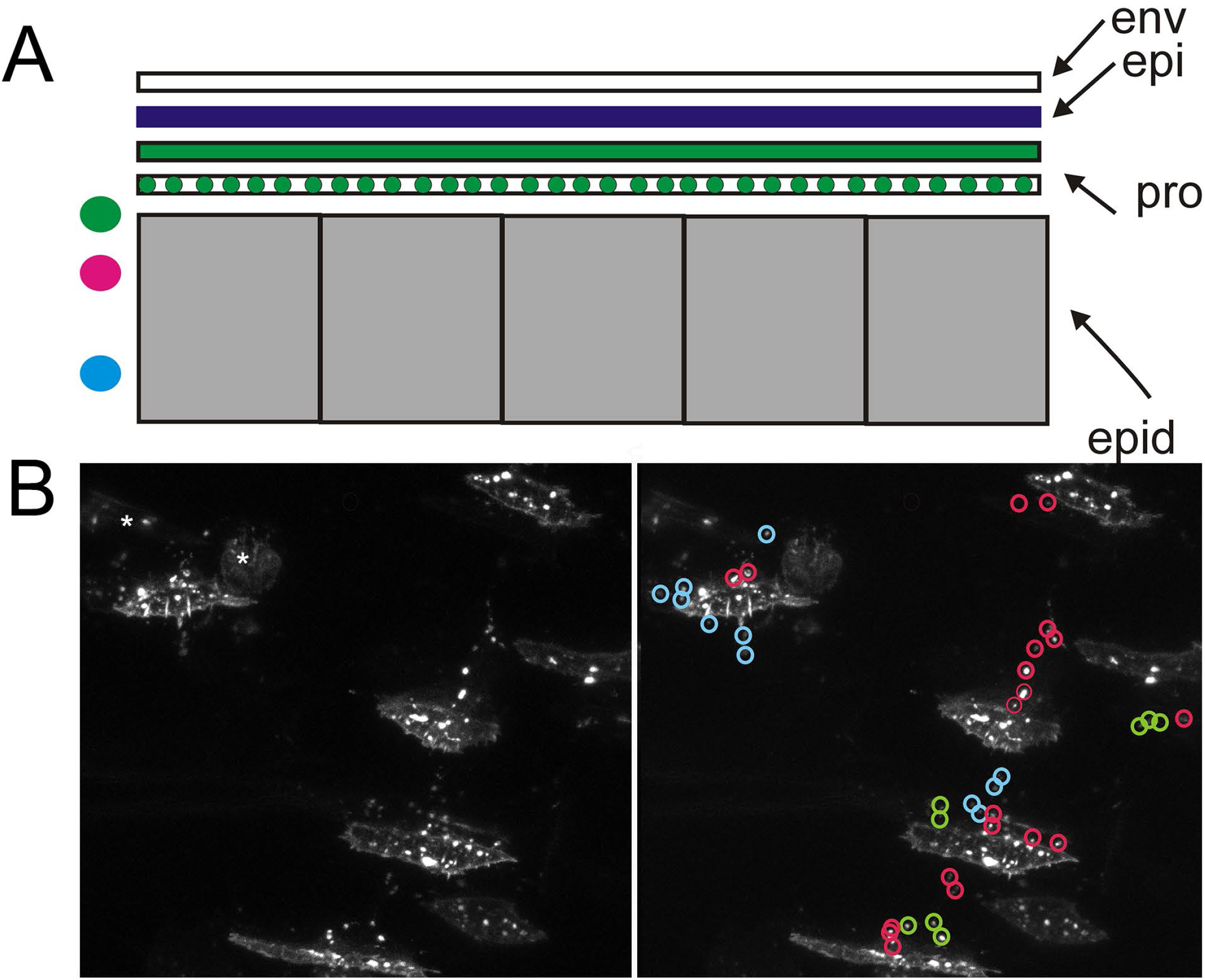
Evidence for the lateral transfer of Kkv::NG. A. A drawing of cuticle during the process of procuticle deposition. The colored circles on the left side show the Z position of puncta from panel B. B. An example of a group of flip out clones in a 70 hr awp abdomen as observed by in vivo confocal microscopy. The panel on the left shows he unmarked image and on the right the puncta that moved laterally from the clones are marked with respect to their Z position. Green for at or above the apical surface. Red for between the apical surface and the middle of the cell and Blue for the lower half of the cell. Some circles encompassed multiple small punta. The cells marked by an asterisk on the unmarked image were not epidermal cells and were located basal to the epidermis.

### TEM examination of pupal cuticle deposition

Another possible mechanism to explain the short distance cell non-autonomy is for a spread of apically secreted chitin. With this in mind we examined thoracic epidermal cells during the synthesis of the pupal cuticle by transmission electron microscopy (TEM). We could identify cell boundaries by the presence of junctional complexes (Figure 8, asterisks). We observed that the undulae found on most epidermal cells during cuticle secretion were often bent over the position of the junctional complex (Fig. 8BEFI). We also often observed what appeared to be “trains” of secreted material between the undualae and the cuticle. These “trains” were often curved and extended over the cell boundary to above the neighboring cell (Fig 8BEHFI). These observations suggest that cuticle material secreted from one cell can end up covering part of a neighbor.

**Figure 8.**
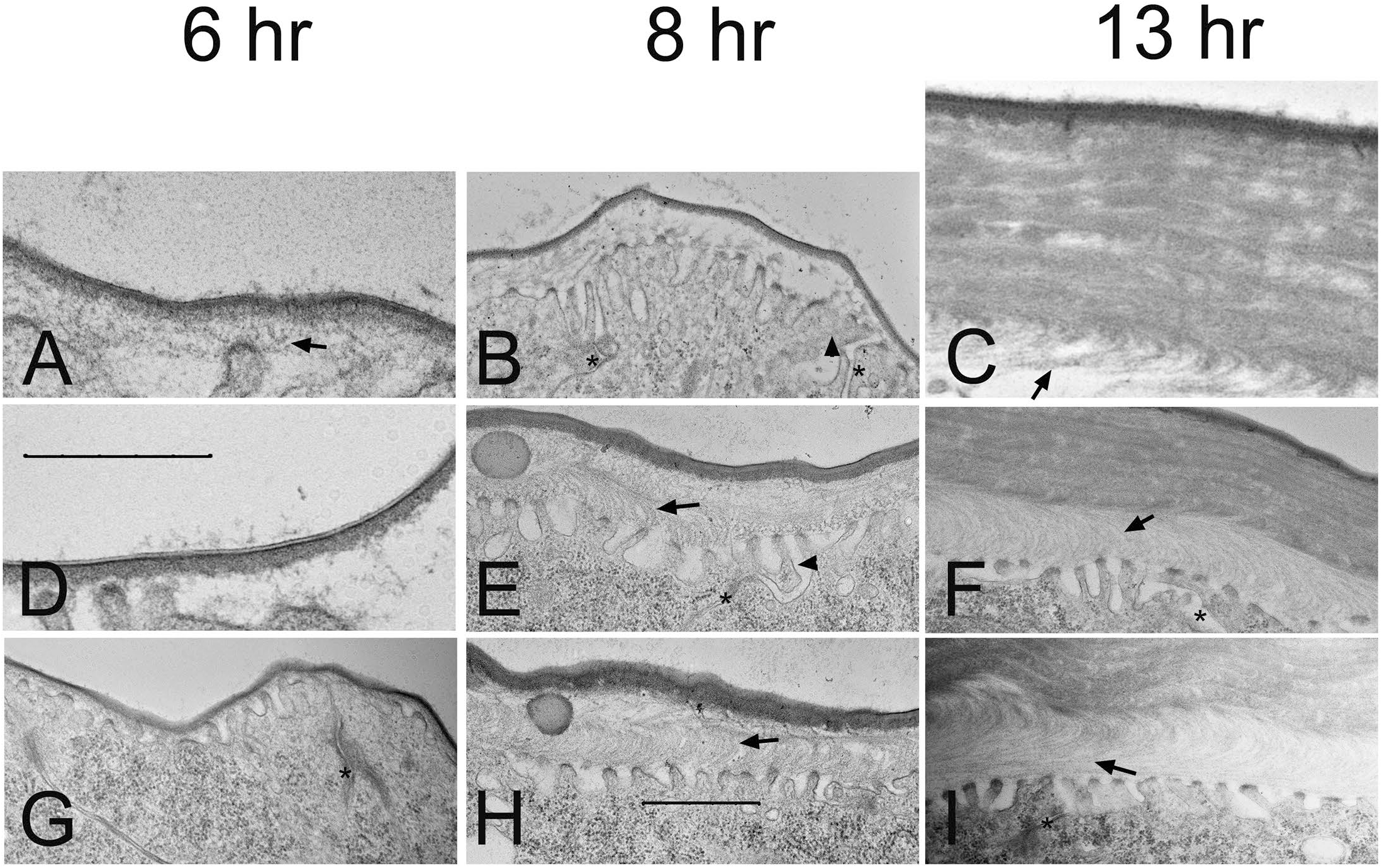
Transmission Electron Microscopy of thoracic pupal cuticle deposition., Panels ADG are of 6hr awp (after white prepupae) animals. Panels BEH are from animals 8 hr awp and panels CFI are from animals 13 hr awp. Arrowheads point to undulate that extend over the junctional complex to be above the neighboring cell. Arrows point to secreted material in “trains” that curve and join the forming cuticle displaced laterally from the undulae it appears to derive from. This is most dramatic in the 13 hr awp animals but is also clear in the 8 hr animals and there are hints in the 6 hr samples. The size markers are 5 um. Panels A and D are shown at a higher magnification than the other images. Panel D is shown to illustrate the distinctive dark/light/dark envelope. Not all micrographs show this as well.

## Discussion

### Multiple mechanisms for timing of the developmental program in pupal wings

The timing of hair growth from wing epidermal cells is not uniform across the wing, as it starts first in distal wing cells and moves proximally in a patchy fashion (Lu et al., 2015; Wong and Adler, 1993). In contrast, cuticle pigmentation, the deposition of chitin and the accumulation of chitin synthase all appear uniform across the wing at different times. Thus, it seems likely that there are at least two different systems functioning in parallel that control the timing of different aspects of the wing developmental program. Is it also worth noting that in other body regions the timing for similar events is different from that seen in the wing. For example, the morphogenesis of hairs and bristles, chitin deposition, chitin synthase accumulation and pigmentation all happen later in abdominal segments. Further, the timing varies from one segment to another with the more posterior segments being delayed compared to the more anterior abdominal segments. The basis for these timing differences remain unknown but are worth investigating in the future.

### Cell non-autonomy of *kkv*

Previous data indicated that *kkv* acted cell autonomously (Ren et al., 2005) but here we established it acted non-autonomously over short distances. The phenotypic data required being able to see the phenotypic changes over part of a cell. The previous experiments were not designed to detect such a subtle phenotype and hence missed it. This short-range non-autonomy is well situated to ensure the continuity of the cuticle across cell boundaries even if neighboring cells are not perfectly synchronized with respect to gene expression. We observed material that appeared to be secreted by undulae moving to the maturing cuticle in a “curved train”. This likely results in cuticle synthesized by one cell contributing to the cuticle that covers part of a neighbor. We also detected the movement of Kkv::NG from clone cells to neighboring cells. This provides a second simple mechanism for short-range cell non-autonomy of *kkv*. Indeed, if a Kkv like movement of transcription factors between neighboring cells takes place it could serve to synchronize gene expression in neighboring cells to promote precise cuticle continuity. I have no evidence on this point however. I note that intercellular transfer between neighboring cells has been seen previously in many systems. For example, it is common in plant cells (Sager and Lee, 2018), is seen for viruses in cultured mammalian cells (Roberts et al., 2015) and intercellular transfer of material through ring canals has been seen in both Drosophila oocytes and in somatic cells (McLean and Cooley, 2013) and also by a non-ring canal mechanism (Li et al., 2015; Yang et al., 2018).

### Robustness of cuticle

The importance of cuticle both as a barrier to the outside and as a support for movement makes its integrity of paramount importance to the success of insects. Flight is a standard feature of insects and it is extremely demanding in terms of energy and structural wear on the cuticle. I previously isolated a hypomorphic allele of *kkv* (*kkv∷smFP)* and here I described several additional phenotypes. One of these was the loss of thoracic macrocheatae. We established that the loss of the bristle shaft increased over time and it appears to be due to the shaft breaking off as the surrounding socket cell remained and appeared normal. The frequency of shaft loss was directly related to the dose of *kkv* as it was substantially more common in *kkv∷smFP/DF* flies than *kkv∷smFP/kkv∷smFP* flies. This argues that having a suboptimal amount of chitin in the shaft cuticle makes it less robust to the wear and tear of fly life.

I also scored the loss of sections of the as wing flies aged. Here the loss of segments started at the margin and often stopped at a vein. Wing veins have been found to block crack propagation in locust wings (Rajabi et al., 2015) consistent with our observations. Prior to carrying out the experiments our thought was that the *kkv∷smFP/Df* wings would be the sensitive to fractures. However, that was not what we observed. Surprisingly *Ore-R* wings were much more likely to lose material than the *kkv∷smFP/Df*. Thus, the wings that seemed likely to have less chitin appeared to be the most resilient. One possible explanation for this is that chitin may function as a stiffening agent and stiffer wings may be more likely to fracture. A second possibility is that *Ore-R* flies may fly faster and put more stress on their wings. The deformation of the thoracic cuticle by indirect flight muscles powers wing beating in Drosophila and many other insects (Pringle, 1949) (Ando and Kanzaki, 2016) (Walker et al., 2014). It is possible that altered physical properties of the thoracic cuticle of *kkv∷smFP/Df* flies due to reduced chitin levels leads to less vigorous wing beating and hence less stress on wings. It would be interesting to test if *kkv∷smFP/Df* flies had different flight characteristics. It would also be interesting to determine if increased *kkv* expression would lead to increased chitin and if that had an effect on fight and wing resilience to fracture.

### Thoughts

There are a number of rather obvious experiments that are not included in the paper. The reasons for this are unusual and worth noting. Several experiments were in progress in early April 2020. These were transmission electron microscopy of *kkv* clones in the abdomen, immune-EM localization of Kkv::NG and examining flipout clones that expressed both Kkv::NG and a red fluorescent membrane tag. The first was to examine in more detail what happens to abdominal cuticle at the juxtaposition of *kkv* mutant cells and nomal cells. The second was to test the hypothesis that Kkv::NG was localized to the undulae or if it was secreted. The commercial high quality anti-NG monoclonal made this experiment quite feasible. The third experiment was to get direct evidence that Kkv::NG derived from a flip out clone was present inside of a neighboring cell. These experiments could not be completed due to the University of Virginia shutting down due to the Covid-19 pandemic of 2020. I was unable to simply put off the experiments and complete them after the pandemic as I retired on May 24, 2020 and my lab was closed. The fly stocks required to carry out these experiments were sent to the Drosophila stock center in Bloomington in early 2020 and I hope that others interested in cuticle synthesis will take advantage of them to do one or more of these experiments.

## Methods and Materials

### Fly Stocks and Genetics

Flies were grown on standard fly food. They were routinely raised at 25°C, but in some experiments, they were raised at 21°C to slow development and in some experiments the flies were grown at 29.5 °C or 29.3 °C to speed up aging. The kkv-RNAi inducing transgenes came from the VDRC (Dietzl et al., 2007) and TRiP collections (Perkins et al., 2015). The experiments described in detail used a TRiP transgene. The VDRC lines were obtained from the VDRC (http://stockcenter.vdrc.at/control/main). The TRiP lines were obtained from the Bloomington Drosophila Stock Center (http://flystocks.bio.indiana.edu/) (NIH P40OD018537) as were many other lines used in the research (e.g. Gal4 lines, Df stocks, *kkv*^*1*^ carrying stock). The *kkv* edited stocks and the various *UAS-kkv* stocks were made by the author in his lab and are described in (Adler, 2020).

### Confocal Microscopy

Immunostaining of fixed pupal epidermal cells during the deposition of cuticle is complicated by the inability of the antibodies to penetrate cuticle after the early stages of its development. Thus, most of the imaging experiments we carried out on Kkv in pupae were done by in vivo imaging of Kkv::NG. In a small number of experiments we examined Kkv-NG in fixed tissue. Inthese experiments we carried out anti-NG immunostaining. Such tissue was only weakly fixed and we did not use animals that were older than around 48 hr after white prepupae (awp). Otherwise, immunostaining of pupal and larval tissues were done as described previously (Nagaraj and Adler, 2012). Imaging of live Kkv::NG containing pupae was done on a Zeiss 780 confocal microscope in the Keck Center for Cellular Imaging. Stained dissected samples were examined on the same microscope.

### Scanning Electron Microscopy

Wings were removed from two day old adult flies. In the experiments described in the paper the wings contained flip out clones (AyGal4) that expressed an RNAi for *kkv* (Trip line – HMC.05880). The hair phenotype overlapped with that seen previously in clones homozygous for *kkv*^1^ with but with a smaller fraction with the strongest phenotype (Ren et al., 2005). The wings were attached to studs with a vertical surface with conducting paint and they were then fractured with a tungsten needle. They were shadowed with platinum and examined in a Zeiss Sigma VP HD field emission Scanning Electron Microscpe (SEM) (NIH 1S10OD011966-01A1) at the University of Virginia Advanced Microscopy Facility.

### Aging experiments

Zero to 1 day old flies of the desired genotypes were collected and routinely 10 females and 5 males were placed in a vial. On some occasions a somewhat lesser number were placed in a vial (e.g. 7 females and 4 males). The smaller number did not affect the results (but see below). In the first experiment, flies were cultured at 29.5 °C and transferred to new vials twice a week (every 3 or 4 Days). On transfer days the flies were anesthetized with CO_2_ and examined for the loss of flies due to death, for the loss of thoracic macrochetae or damaged wings. In the second experiment, flies were cultured at 29.3 °C and transferred to new food every 3 days. The flies were scored for the same 3 phenotypes every 5 days. When a fly showed a new phenotype between 2 time points the phenotype was estimated to have arisen at the mid-point between the two scoring days. Only data for females are reported in the ms.

We also carried out an experiment where 1 male and 1 female were cultured in a vial with transfer every 3 days. Every 5 days the flies were anesthetized with CO_2_ and examined for the same 3 phenotypes. Micrographs of the anesthetized flies were obtained using an Infiniprobe TS-160 (Infinity Photo-Optical Co) connected to a Nikon Z50 camera. In these experiments it appeared that the loss of bristles was delayed due to the smaller number of flies in the vial however the number of flies examined was too small to be confident of the result.

## Supporting information

all supplemental files

## Acknowledgements

This research was supported by funds provided by the W. R. Kenan Chair to the author and to a limited extent personal funds of the author. The author thanks H.S. Tzu for helpful conversations. “We acquired confocal images using the Keck Center Zeiss 780 Confocal microscopy system (NIH OD016446). We acquired Scanning Electron Microscope images at the Advanced Microscopy Facility at the University of Virginia. The images were obtained on a Zeiss VP HD SEM field purchased with a grant from the NIH (NIH 1S10OD011966).

