## Supplementary material for "Short distance non-autonomy and intercellular transfer of chitin synthase in Drosophila": all supplemental files

**Supplemental Figures**

Figure. S1. Pigmentation in a pupae is similar for neighboring cells. A. All images are of lightly fixed 72 hr pupal tissue by light microscopy. A. Wing. B. Notum. C. Head. The similar level of pigmentation of neighboring hairs in A and of bristles in B and C.

Figure S2. Life span of different genotypes. Data for the two experiments described in the methods are shown. P values are for T tests where we compared the observed life spans to that of other genotypes.

Figure S3. Progressive loss of wing material with aging. The data for the two experiments described in the methods is shown. The P values are from chi square tests with the observed frequency for wing defects for *Ore-R* (or other genotypes) used to generate expected number to compare to the observed frequencies. Panels A-F show examples of wing defects observed in similarly cultured flies. Asterisks mark locations of loss of wing material. Note that in some cases it is clear that there were multiple independent locations of wing margin loss (e.g. Panel D). The wings in ABC are from *kkv::smFP* flies and the wings in DEF are from *Ore-R* flies.

Figure S4. Clones of *kkv* in the abdomen do not show sharp borders of pigmentation. A and B show the the naturally sharp pigmentation borders. C-J show *Ay-Gal4 kkv-RNAi* flip out clones. Note the uneven borders in all of the clones.

Figure S5. Use of SEM for examining procuticle layering. A fracture segment of adult abdominal cuticle imaged on a scanning electron microscope.

Fig S6. Kkv::NG dependent puncta in live pupae. A. Many puncta (arrow) are visible in the space between the pupal cuticle and the notum epidermis in 50 hr awp pupa. The pupal cuticle is visible (asterisk) due to autoflourescence. B. Puncta are visible and concentrated along the mid line of a 20 hr awp *kkv::NG* pupa. At this time the epidermal cells are still attached to the pupal cuticle they synthesized so the cellular outlines are visible. C. Puncta are visible (arrow) in the space between the pupal cuticle and the notum epidermis in 50 hr awp *ap-Gal4/+; UAS-kkv::NG/+* pupa. D. No puncta are visible (arrow) in the space between the pupal cuticle and the notum epidermis in 50 hr awp *ap-Gal4/+;*

*UAS-kkv R896K::NG/+* pupa. This is likely due primarily to the protein not being localized properly to the apical surface.

Figure S7. Neon Green is a useful reporter for Kkv::NG in secreted puncta. Pupal cuticle was fixed and dissected from 42 hr *kkv::NG* pupae and immunostained with both anti-NG (green) and anti-Kkv (red) antibodies and imaged by confocal microscopy. Many puncta were visible and a large number of these stained with both antibodies arguing that NeonGreen is a useful reporter for Kkv in cuticle. Note the large number of co-labelled (yellow) puncta (e.g. arrows).

Figure S8. Pupal cuticle has a different structure in different regions of the animal. In vivo imaging of *kkv::NG* pupal cuticle. The morphologies of the thoracic (TH) and abdominal (Abd) pupal cuticles are different. Similar differences are also seen with *Ore-R* pupal cuticle so this represents a difference in autofluorescence. The arrow points to several Kkv::NG puncta. These are not seen in *Ore-R* pupae.

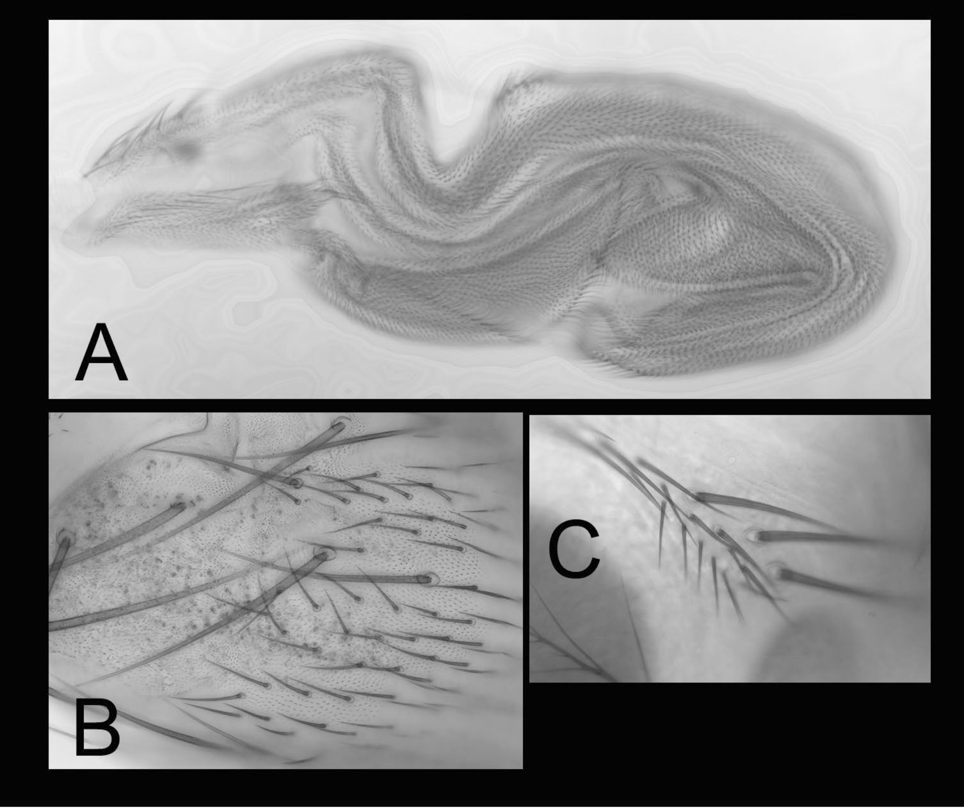


Figure S1

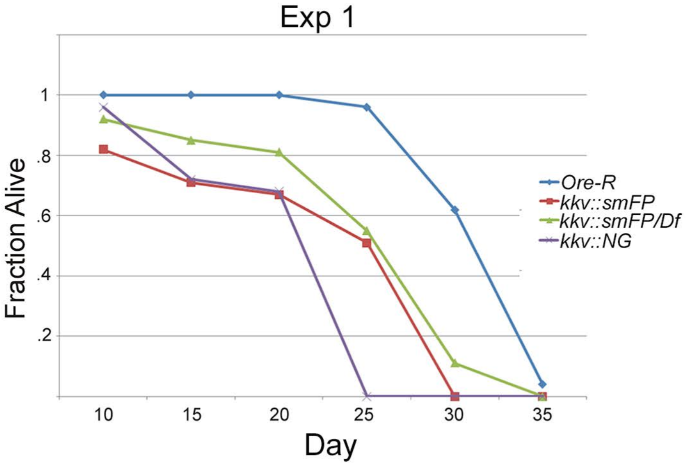


| | *Ore-R* | *kkv::smFP* | *kkv::smFP/DF* | *kkv::NG* |
|---|---|---|---|---|
| number of flies | 50 | 51 | 62 | 25 |
| mean | 25.28 | 20.27 | 23.32 | 18.7 |
| SD | 2.78 | 8.62 | 7.12 | 4.04 |
| P compared to *Ore-R* | NR | 0.00021 | 0.06966 | 1.37E-08 |
| P compared to kkv::smFP | 0.00021 | NR | 0.053 | 0.25 |

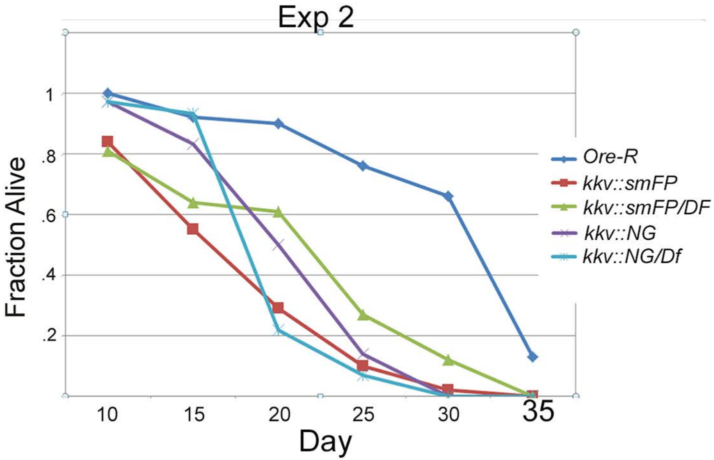


| | *Ore-R* | *kkv::smFP* | *kkv::smFP/* | *kkv::NG* | *kkv::NG/D* |
|---|---|---|---|---|---|
| number of flies | 38 | 51 | 59 | 36 | 73 |
| mean | 29.34 | 16.18 | 19.79 | 19.71 | 18.46 |
| SD | 6.92 | 6.07 | 8.32 | 5.01 | 3.69 |
| P compared to *Ore-R* | NR | 3.67E-15 | 5.96E-08 | 2.3E-09 | 1.92E-19 |
| P compared to *kkv::smFP* | 3.67E-15 | NR | 0.012 | 0.0053 | 0.011 |
| P compared to *kkv::NG* | 2.3E-09 | 0.0053 | 0.95 | NR | 0.19 |

### Figure S2

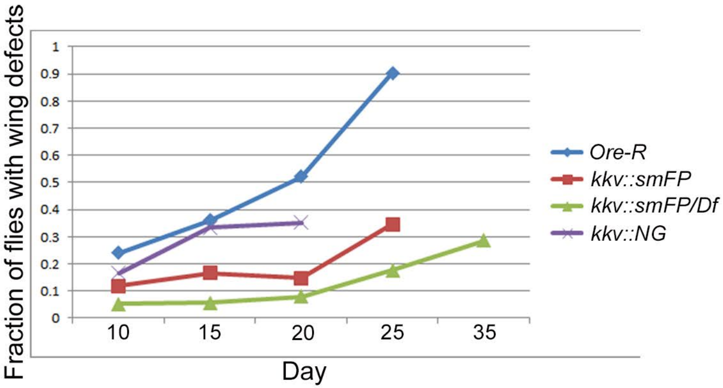


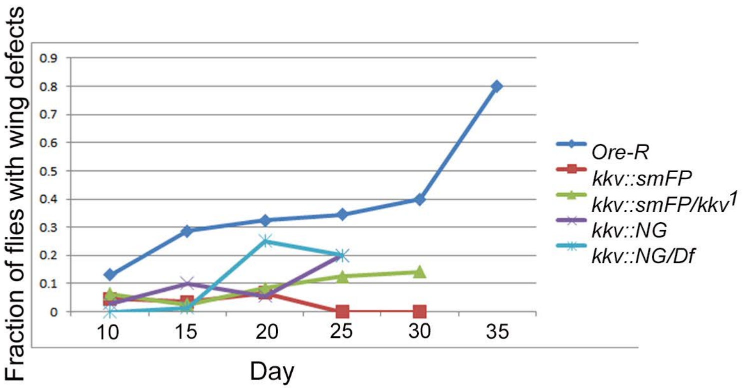


| | *Ore-R* | *kkv::smFP* | *kkv::smFP/* | *kkv::NG* |
|---|---|---|---|---|
| p vs Ore-R | | **0.0000128** | **4.31E-10** | 0.16 |
| *p vs kkv::smFP* | **2.65E-13** | NR | 0.18 | **0.013** |

| | *Ore-R* | *kkv::smFP* | *kkv::smFP/kkv*[1] | *kkv::NG* | *kkv::NG/Df* |
|---|---|---|---|---|---|
| p vs *Ore-R* | NR | **0.035** | **0.0023** | **0.016** | 0.55 |
| p vs *kkv::smFP* | **2E-09** | NR | 0.69 | 0.85 | **0.0037** |
| p vs *kkv::NG* | 9.18E-12 | 0.85 | 0.47 | NR | **0.00069** |

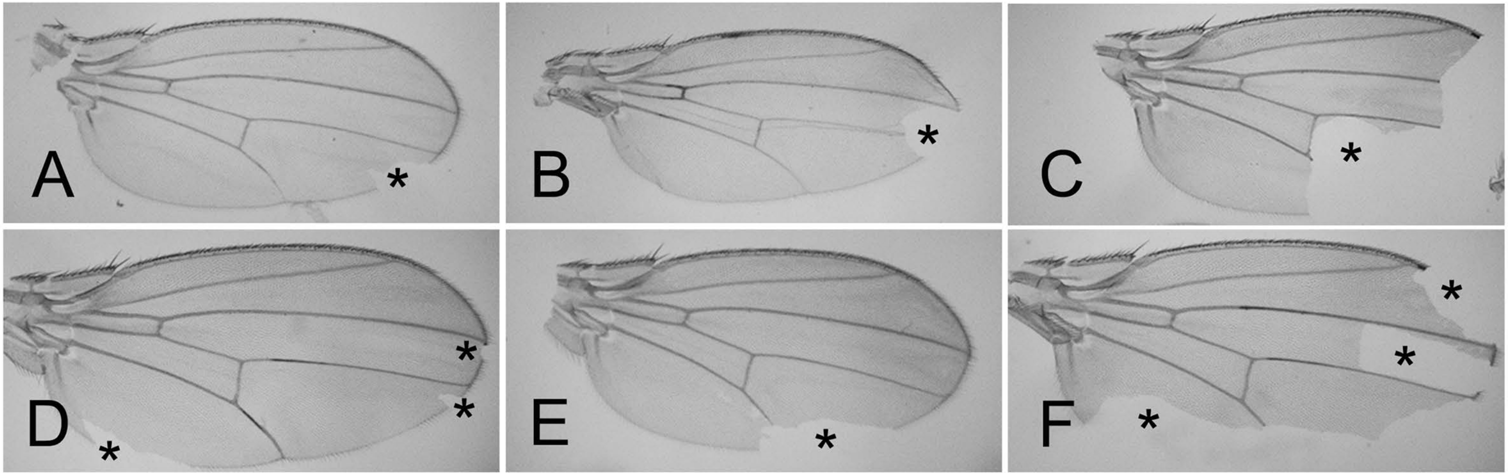


Figure S3

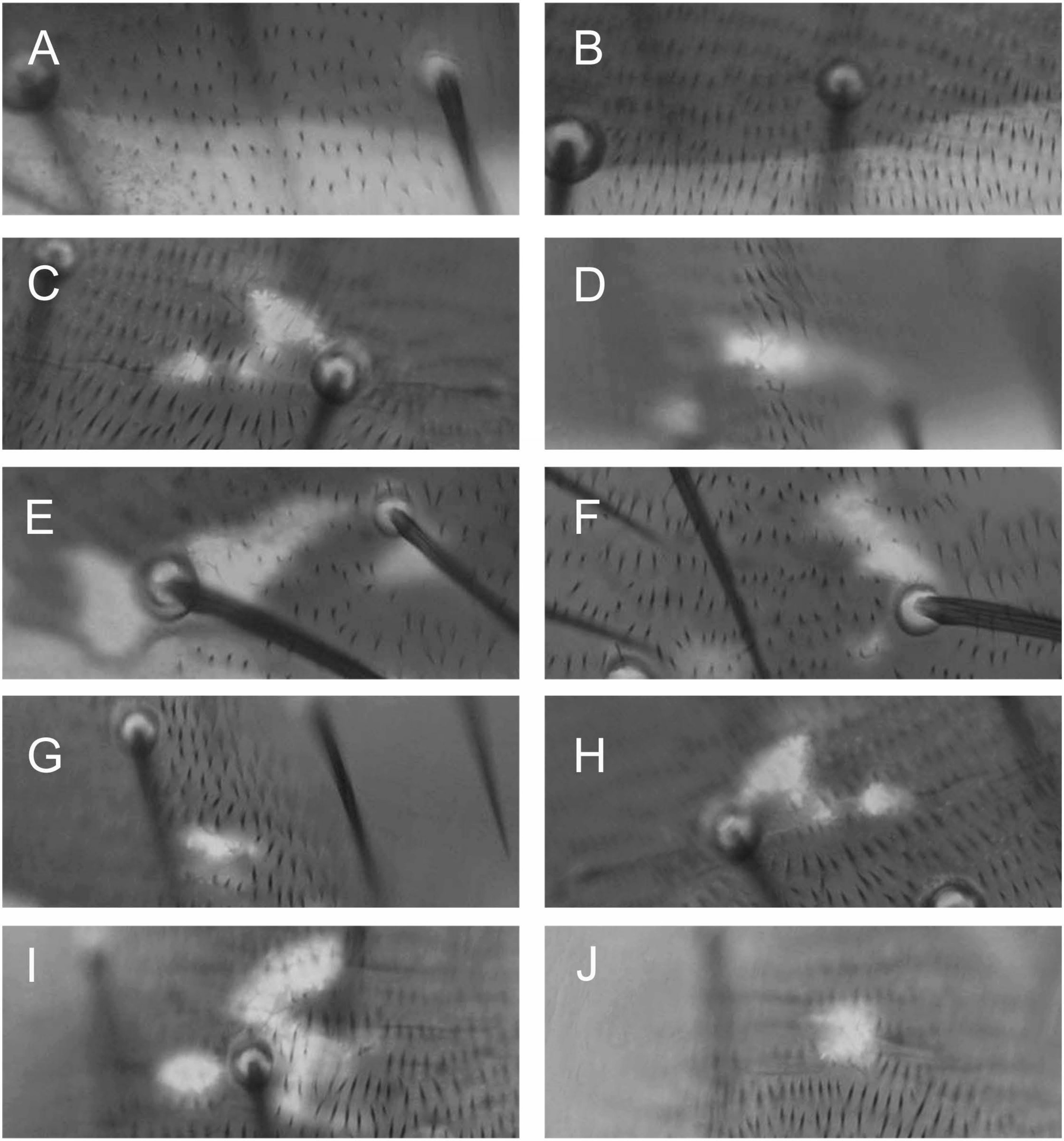


Figure S4

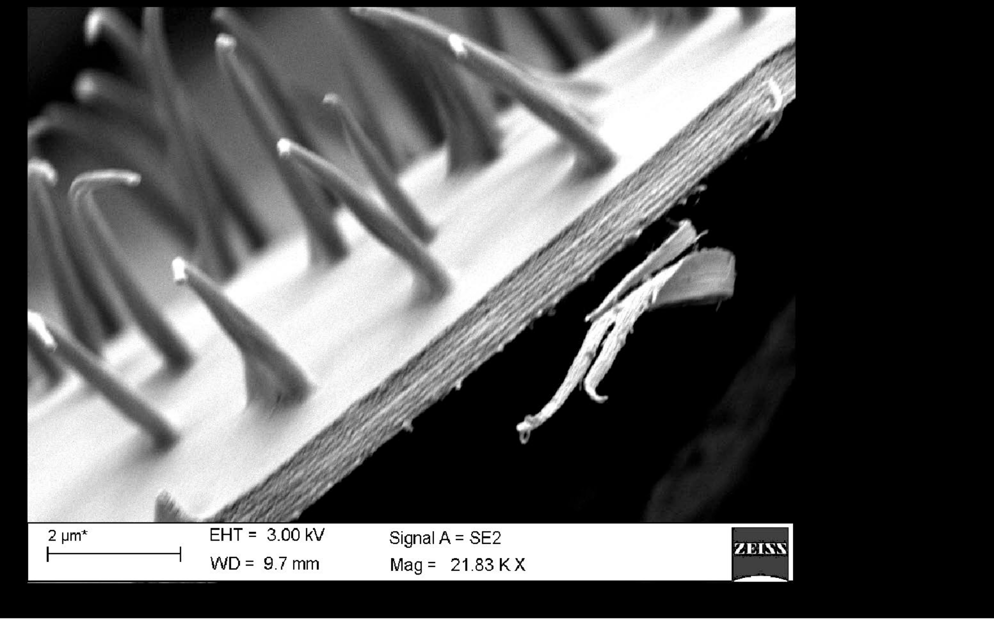


### Figure S5

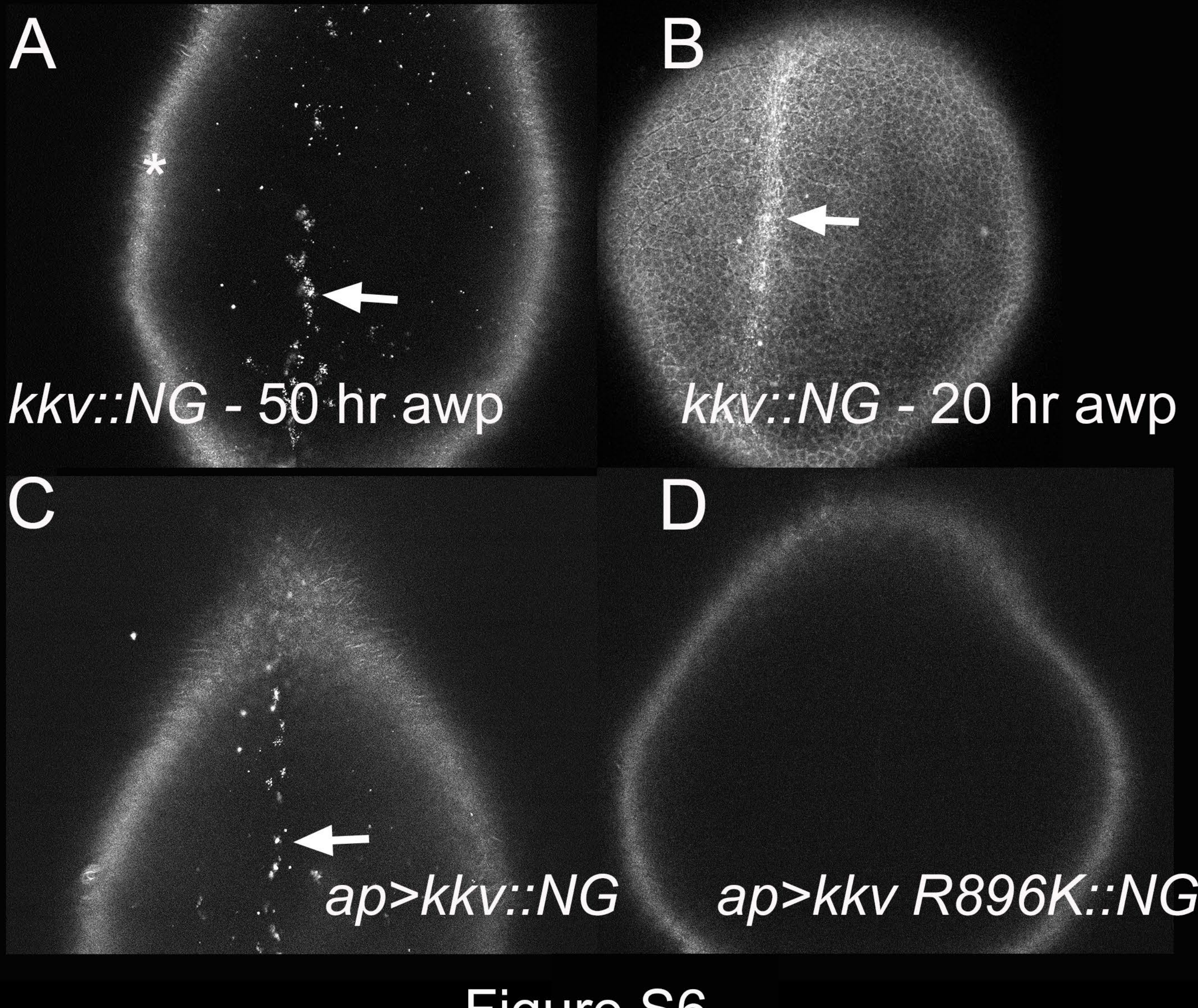


Figure S6

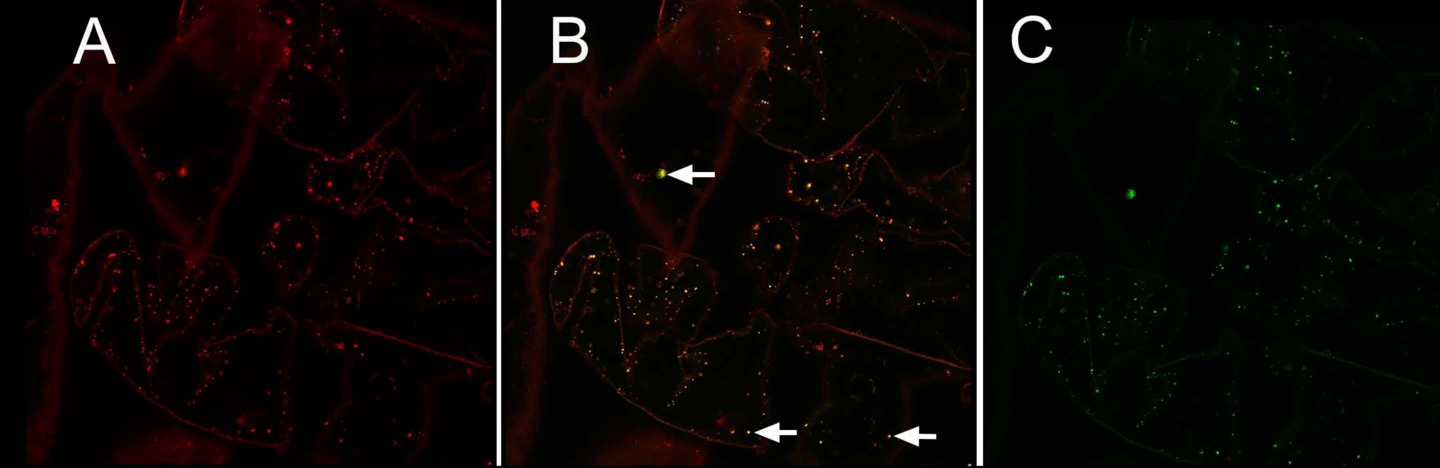


Figure S7

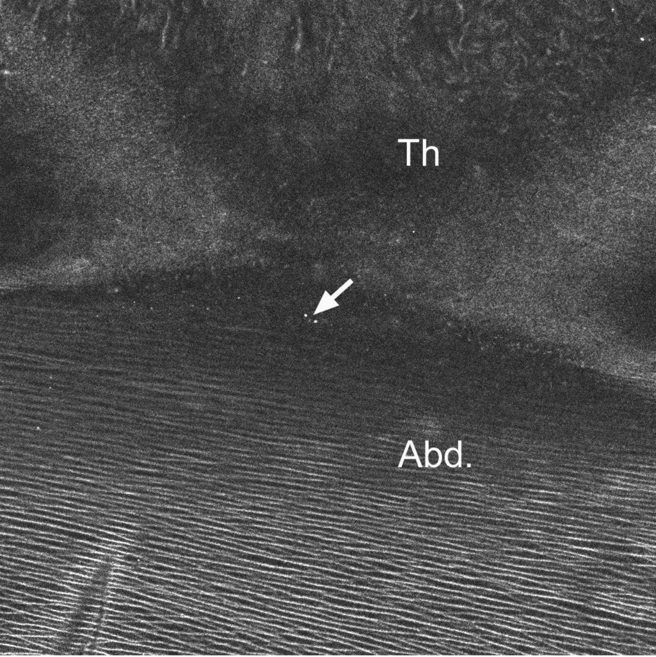


Figure S8
